# A minimalist binary/digital approach to large-scale single molecule protein identification with optically labeled tRNAs and multiple carboxypeptidases and its extension to peptide sequencing

**DOI:** 10.1101/2024.12.02.626402

**Authors:** G. Sampath

## Abstract

Recently a binary/digital scheme based on the superspecificity property of transfer RNAs (tRNAs) was proposed for the identification of single amino acids (AAs) from binary-valued measurements (*Eur. Phys. J. E* **45**, 94, 2022). There are two formulations, they can be used to sequence short peptides and/or identify their parent proteins. In one of them an array of peptides is sequenced in 20 cycles by adding 20 different tRNAs carrying a fluorescent tag, optically recognizing the C-terminal residues, and cleaving the latter with a carboxypeptidase; the process is repeated over the peptides in parallel. Here this scheme is used to develop in theory a minimalist approach to protein identification that uses only two tRNAs and the carboxypeptidases A, B, and C. The latter form a complete and mutually exclusive set capable of cleaving all 20 AA types; this divides the 20 AAs into three classes. The sequences obtained are partial sequences in the reduced alphabet, their parent proteins can be obtained by search through a proteome database. The AA class of the terminal residue of every peptide in the array can be identified in a single cycle by using the three carboxypeptidases in the order C-B-A. With peptide lengths of ∼20 and a cycle time of ∼1 hour, the parent proteins of K peptides can be obtained in about 20 hours. This is independent of K (within the limits imposed by the imaging method used) and the dynamic range of a proteome; thus in theory a whole proteome can be processed in less than a day. Computational results suggest that the parent proteins of over 92% of peptides from the human proteome (Uniprot id UP000005640_9606) can be identified. The identification rate when residues are skipped due to carboxypeptidases cleaving the second and later residues in delayed reactions is about ∼90% with 1 or 2 skips. Full sequencing without skipped residues can be done by using all 20 tRNA types over 20 cycles in increasing order of cleavage time of the 20 AA types; a recursive procedure is given. A procedure for AA identification in the bulk with a minimal number of steps that can be implemented with equipment in an undergraduate lab is also included; it may be suitable for a range of projects at the junior/senior level.

## 1. Protein sequencing and identification in the bulk and with single molecules

Protein sequencing has for several decades been done with one of three methods: Edman degradation, gel electrophoresis, and mass spectrometry (MS) [1-3]. All three are bulk methods that require samples typically containing, at a minimum, several million copies of a protein. Although with continually improving purification and separation techniques significantly lower sample sizes are becoming possible, MS is yet to solve the dynamic range problem [4] posed by vastly different copy numbers of the constituent proteins of single cells, whose number may vary from a few tens of copies of one protein to millions of copies of another. Random sampling of the assay sample to reduce the number of molecules examined is invariably biased toward the more populous proteins. To reduce sampling bias the assay sample is selectively separated into fractions such that proteins with smaller copy numbers have a better chance of getting into the spectrometer. These difficulties have led to the development of single molecule protein sequencing (SMPS) and identification methods designed to work with single molecules or small numbers (∼10-100) of them [5,6]. Some of them are modifications of techniques used in next generation sequencing of DNA. SMPS methods include: nanopore-based methods [7]; plasmonics [8]; methods based on tethered arrays of peptides similar to microarrays, with selected residues carrying fluorescent tags [9] or with antibodies that bind with short sequences of amino acids [10]; and semiconductor chips containing arrays of wells [11] where the N-terminal residues of peptides are allowed to bind preferentially with specially designed N-terminal amino acid binders (NAABs) added to the wells.

### Sequencing and identification in the bulk

Protein sequencing of bulk samples is largely based on Edman degradation (ED), in which the N-terminal residue of a peptide is targeted for removal and derivatized in the process. The derivatized cleaved residue is identified with spectrographic techniques. ED has been the staple for almost seven decades as it can be used to reliably cleave residues in sequence order from the N-terminal. However it is a harsh chemical-based method as it degrades the cleaved residue and is also time consuming. Even with automated procedures cleaving and identifying a single terminal residue requires about 10 minutes. For this and other reasons ED is limited to proteins with about 50-70 residues. Several other less harsh methods based on chemical reagents have been devised, for example [12]; almost all of them are based on derivatization followed by spectrographic identification of the derivatives.

This has led to the development of non-chemical alternatives, including the use of enzymatic methods in which exopeptidases cleave the N-terminal or C-terminal residue while leaving it largely intact. Aminopeptidases, which are exopeptidases that cleave the N-terminal residue, have been studied in detail. In [13], leucine aminopeptidase has been investigated, it is capable of cleaving a majority of the AAs. A number of carboxypeptidases, which are exopeptidases that cleave the C-terminal residue, are available [14]. Serine carboxypeptidases have been found to be effective. In particular carboxypeptidase Y (CPD-Y) and malt carboxypeptidase (CPD M-II) have been found to be fairly reliable. Specificity and cleavage interval times vary widely and depend on solution pH and the substrate. Thus CPD-Y preferentially cleaves hydrophobic residues, CPD-M-II does the same with basic residues. Other well-studied carboxypeptidases include carboxypeptidase A, carboxypeptidase B, and carboxypeptidase C [15], these three are the subject of the present study as they constitute a complete set of mutually exclusive subsets capable of cleaving all 20 AAs. Protein identification in the bulk is most commonly done with antibodies, which implies prior knowledge of the proteins present in a sample [16].

### Single molecule sequencing and identification

With increasing evidence of the role played by sparsely occurring proteins in single cells in disease conditions there has been a shift toward single cell protein sequencing. Given the small sizes of the samples obtained and the dynamic range of copy numbers alluded to earlier bulk methods are not suited to the task. This has provided additional motivation for SMPS methods, some of which are based on cleaving of terminal residues and use ED or exopeptidases. For example, in [9] optical measurements of fluorescence levels due to tagged residues in a peptide array are followed by a cleaving step in which ED is applied to the tethered ensemble. In [11], aminopeptidases are used to cleave the N-terminal residue after optical measurements of binding times for the residue with different NAABs have been made. AA-specific enzymes collectively named Edmanase [17] have been designed expressly for use in protein sequencing. Nanopore-based methods are among the most widely studied methods. Nanopore protein sequencing is reviewed in [18]. Recently reported methods include [19,20]. In [21] an electrolytic cell with two nanopores in tandem has been proposed. In this method, which has not been implemented, the leading terminal of a peptide is cleaved by an exopeptidase covalently attached to the cell membrane on the exit side of the first pore. The cleaved residue then translocates through the second pore where it is identified based on the volume excluded and its dwell time in the pore. Several studies have focused on identifying single AAs [22-24] or small classes that AAs belong to [25]..

### The present study

The potential use of carboxypeptidase A, carboxypeptidase B, and carboxypeptidase C as a complete set capable of cleaving all 20 AAs at the single molecule level in the binary-digital method of [24] is at the center of this study. The C-terminal residues in an array of peptides immobilized at their N-ends on a glass slide can be identified in parallel with a set of cognate tRNAs, following which the three carboxypeptidases are applied n the order C-B-A to cleave them. The partial sequences so obtained can be used in conjunction with a proteome database to identify their container proteins in parallel. For most peptides of length 20-50 (residues) this can be done in a reasonable amount of time [26]. This expectation is supported by the results of computational experiments on peptides obtained from a random set of proteins in the human proteome (Uniprot id UP000005640_9606); the results show that 90-92% of the parent proteins of all peptides from the human proteome can be identified unambiguously under certain conditions.

A procedure to implement the above AA identification method in the bulk using minimal steps and equipment commonly found in an undergraduate chemistry lab is outlined; its potential use for experiments and projects at the junior or senior level is indicated.

## 2. A binary/digital method for identifying the terminal residues of peptides at the single molecule level

Almost all of the protein sequencing/identification methods alluded to in Section 1, whether in the bulk or for single molecules, are based on precise measurement of an analog quantity (blockade current, mass, electrical charge, binding time, fluorescence intensity level, etc.). The method proposed in [24] moves away from analog measurements to a binary/digital approach based on the ‘superspecificity’ property of tRNAs. Thus for every AA there are one or more tRNAs that bind only to that AA; superspecificity ensures a very low error rate (less than 1 in 150; with built-in *in vivo* error correction the error rate is much lower). The proposed method can be used with bulk samples as well as single molecules (or small numbers of them). This is a *de novo* method that does not require *a priori* information about the protein/peptide or a database for comparison.

There are two formulations. In the first an AA is identified in four steps: confinement, filtration/separation, deacylation, and detection; it is executed in parallel on 20 or more copies of AA each with a different tRNA. AA detection may be optical based on fluorescent tags (a single color is sufficient, actually any color can be used for any of the AAs); it uses only 3 of the 4 steps (the deacylation step is not required). Alternatively detection can be electrical based on translocating the freed AA (after it has been dissociated from its cognate tRNA) through a nanopore and measuring the change in the current, it requires all 4 steps.

Figure 1 summarizes this first approach. See [24] for the details.

**Figure 1.**
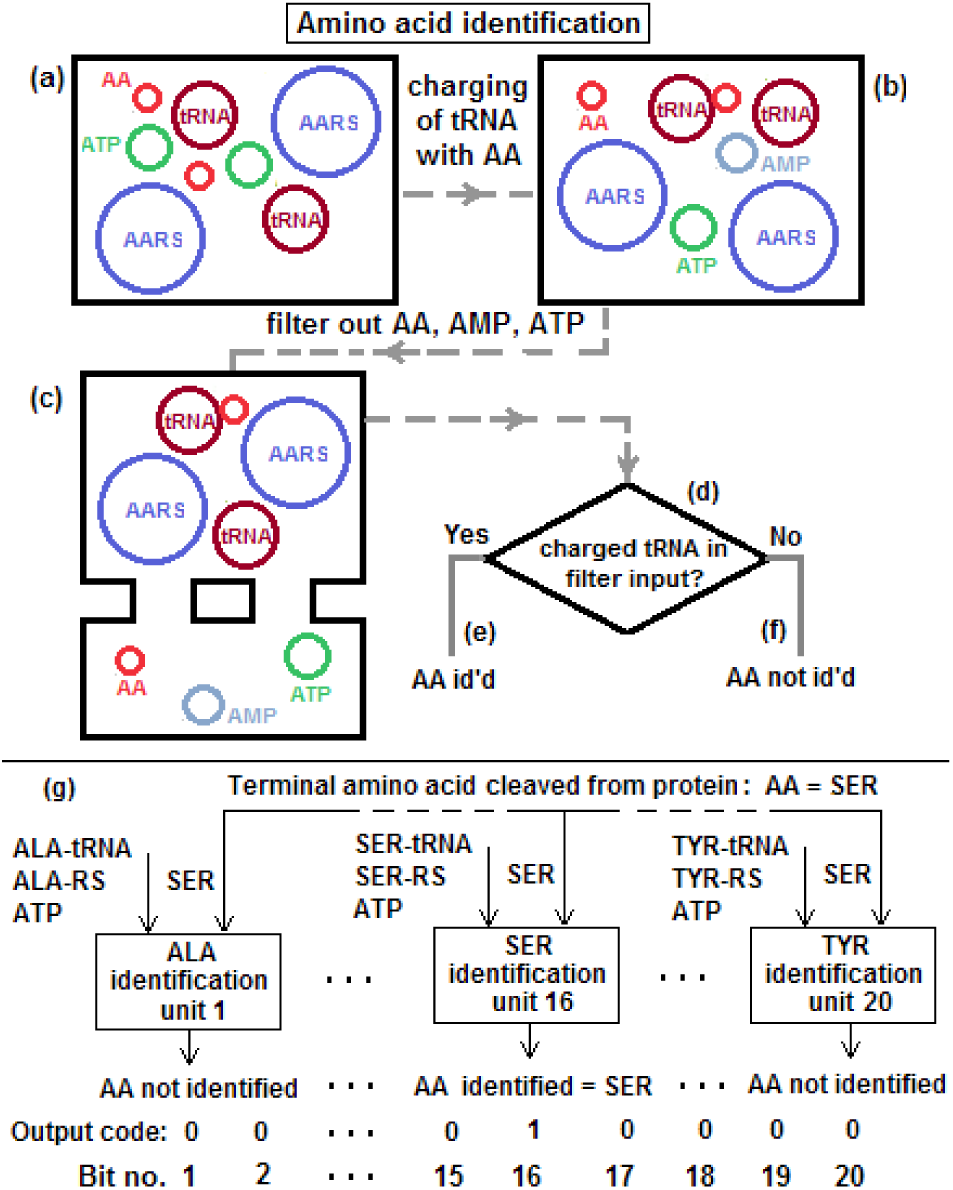
Binary/digital identification of free amino acids (AA) obtained by cleaving terminal residue at N- or C-end of peptides in purified sample. Parts a and b show schematic of binary AA detection unit. a, b: Fluorescent tag attached to free AA, added to cavity with one type of tRNA, its cognate AARS, and ATP; tRNA gets charged with cognate AA. c: all smaller molecules (excess ATP, AMP (if charging occurs), unbound AAs) removed by filtration. d, e, f: presence of charged tRNA detected optically with total internal reflection fluorescence (TIRF); or electrically: cognate tRNA dissociated with NaOH, freed AA passed through nanopore in electrolytic cell, its presence detected from occurrence of current blockade. g: 20 AA detection units, one per AA type, 20-bit digital output corresponding to 20 copies of AA input has one bit high, all others low. (Figure from [24].)

A second and somewhat less complex approach that does not require purification uses superspecificity for parallel sequencing of large numbers of peptides bound to a glass slide. In 20 cycles 20 different tRNAs are used to identify the terminal residues of the peptides in the array, following which a carboxypeptidase is used to cleave the detected terminal residues across the array.. This approach can be viewed as a binary-digital alternative to the analog methods of [9] and [11]; it requires only a single parallel image measurement of the peptide array per terminal residue across the array.

Figure 2 summarizes this second approach; see [24] for the details.

**Figure 2.**
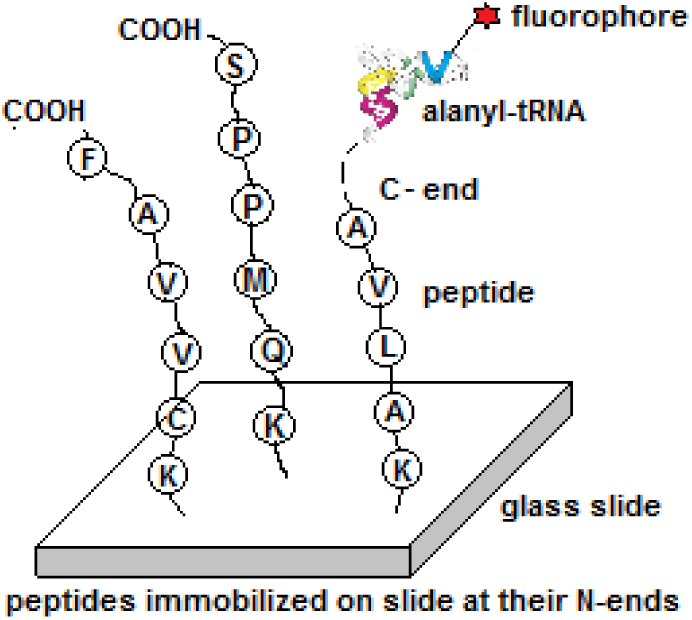
Sample peptides from mixture bound at their N-ends to glass slide microarray. In 20 passes, 1 ≤ i ≤ 20, tRNA_i_, AARS_i_, and ATP molecules added to solution; tRNA_i_s bind with all cognate terminal residues AA_i_ at C-end; TIRF image captures locations of charged tRNA_i_s in microarray. Following deacylation, freed tRNA_i_, cognate AARS_i_, and excess ATP molecules washed out. Process repeated until all peptides have been sequenced. (Figure from [24].)

Following [24], parallel sequencing of an array of peptides immobilized at their N-ends on a glass slide can then be done as follows:

**Procedure** SequencePeptideArray

1. Bind mixture of peptides at their N-ends to glass slide. Initialize sequence data array **S** to empty.
2. Add fluorescent tags to set of N = 20 unique tRNAs. (Single color sufficient, can be used for all 20 tRNA types (because two different tRNAs cannot be simultaneously present).
3. For i = 1 to N do the following:
  3a: Add tRNA_i_ AARS_i_, and ATP to solution. If C-terminal residue of a peptide is cognate to tRNA_i_ latter gets charged with that residue.
  3b: Wash away excess reactants.
  3c: Image peptide array with TIRF. Only those array positions corresponding to peptides with cognate terminal residues will have high signal (that is, different from background signal). Add i to corresponding sequences in sequence array **S**: i identifies next residue in sequence for peptide.
  3d: Deacylate charged tRNAs with NaOH. This returns peptide array to starting state before Step 3a with one less teminal residue for all peptides with terminal residue = AA_i_.
  3e: Wash away uncharged tRNAs, AARS, excess ATP, AMP (if tRNA_i_ is charged).
4. After Step 3 every terminal residue in every peptide has been identified.

Cleave all terminal residues with a carboxy peptidase to expose next residue in each peptide in peptide array.

Repeat Steps 3 and 4 until fluorescent signal = background signal.

In the following sections this second formulation is reduced to a minimalist form in the context of terminal residue cleavage based on three different carboxypeptidases: A, B, and C. By using only two tRNA types and applying the three peptidases in the appropriate order fairly accurate sequencing of large numbers of bound peptides may be done simultaneously. The errors that occur because of delayed reactions between the carboxypeptidases and the substrate array are also considered. It is shown that even with these errors present the partial sequences obtained can in general be used to simultaneously identify the parent proteins of all the peptides in the array. In addition to the digital nature of the measurements (which makes the sequencing/identification process more robust than analog measurements), scaling from a single peptide to very large numbers of them makes the method a viable alternative to several large-scale methods that are currently being developed or translated into lab or field use [9-11].

## 3. C-terminal residue cleavage with multiple carboxypeptidase types

Terminal residue cleavage with carboxypeptidases is reviewed in [15]. Briefly there are two ways to identify the C-terminal residue of a peptide. The first is based on the use of hydrazine, which forms derivatives of all the residues in the peptide except the C-terminal residue. Spectrographic methods can then be used to identify the remaining C-terminal residue. This is somewhat wasteful as the entire peptide is used to identify just one residue, so it is rarely used. The second way is to cleave the C-terminal residue successively from the peptide and identify it by using spectrographic or other methods. Generally these approaches are for bulk samples. As noted earlier, with single cell analysis gaining importance residue identification must be able to deal with very small samples, down to a few tens of peptide molecules. Spectrographic methods are not useful here as they are inherently designed for sample sizes from micrograms down to at best a few nanograms perhaps, which means at least 10^12^ molecules in the sample.

Some of the exopeptidase-based methods mentioned above have recently been adapted to single molecule analysis. These efforts use Edman degradation or an aminopeptidase to cleave the N-terminal residues of a massive array of peptides [9,11], thus gaining in volume what is lost in time. In contrast very few single molecule methods based on the use of carboxypeptidases for C-terminal residue cleavage, other than, perhaps, the second formulation in [24] referred to above, have been reported. The present work looks at how such usage can result in reasonably successful sequencing of large numbers of peptides and/or protein identification. The latter is based, ironically, on the fact that the sequencing is error-prone yet yields partial sequences that can be used to effectively identify the parent proteins of the partially sequenced peptides. The following property is central to this study:

### Carboxypeptidase A, B, and C form a complete and mutually exclusive set of exopeptidases for protein sequencing

Thus carboxypeptidase A, B, and C are together able to cleave residues of all 20 AA types: A cleaves all C-terminal AA types except Lys, Arg, and Pro; B cleaves Arg and Lys; C cleaves Pro. Thus when used collectively (in some order), all 20 types of C-terminal residues in an array of peptides can be cleaved. Cleaving the terminal residue across an array of peptides can then be done in a single cycle of 3 steps. In an ideal situation every peptide has its terminal residue cleaved in the same one of three steps. The optimal order is:

* Cleave all Pro terminal residues with carboxypeptidase C
* Cleave all Arg and Lys terminal residues with carboxypeptidase B
* Cleave terminal residues of the other 17 AA types with carboxypeptidase A.

Assuming that the peptides are obtained by proteolysis resulting in strings with Lys at the N-end (and nowhere else), the sequencing process requires M cycles, where M is the largest value for the peptide length across the array. Procedure SequencePeptideArray can be rewritten as follows:

**Procedure** SequencePeptideArrayWithCarboxypeptidaseABC

1. Bind mixture of peptides with Lys at their N-ends to glass slide. Initialize sequence data array **S** to empty.
2. Add fluorescent tags to tRNA_Pro_ and tRNA_Arg_. A single color is sufficient if recognition/cleavage of Pro and Arg is done in series; two colors need to be used for recognition-cleavage in parallel, with both tRNAs added together. (Notice that the other 17 AA types do not have to be tagged, as they are all recognized by default as the AA type X; see Section 4 below on the resulting 3-character alphabet {P, R, X}.)
3. Recognition-cleavage:
  3a: If C-terminal residue of a peptide is cognate to tRNA_P_ or tRNA_R_ the latter gets charged with Pro or Arg.
  3b: Wash away excess reactants.
  3c: Image peptide array with TIRF. Only those array positions corresponding to peptides with cognate terminal residues will have high signal (that is, different from background signal). Add P or R to corresponding sequences in sequence array **S**: i identifies next residue in sequence for peptide. Add X to every other sequence.
  3d: Deacylate charged tRNAs with NaOH. This returns peptide array to starting state before Step 3a.
  3e: Wash away uncharged tRNAs, AARS, excess ATP, AMP (if tRNA_P_ or tRNA_R_ has been charged).
4. After Step 3 every terminal residue in every peptide has been identified as being P, R or X.

Add carboxy peptidase C and B if recognition-cleavage is in parallel or C followed by P if done in series.

Add carboxy peptidase A to cleave all the X terminals.

This exposes next residue across peptide array.

Repeat Steps 3 and 4 until fluorescent signal = background signal.

In practice cleavage of a terminal residue will not be perfect over time because the following residues (second, third, etc.) may also be cleaved if the reaction continues. It is therefore necessary to determine when to stop the reaction. Empirical studies can be done for each of the 20 AA types to obtain their reaction times (see end of Item 1 in Section 8). Even with this information, with an array of peptides it is not possible to stop the reaction exactly after each terminal residue is released because the exact times may be different for each peptide, whose terminal residue could be different from that of the others in the array. However, while this has the potential to increase sequencing errors across the array, it does not have as adverse an impact on protein identification. This is considered in the next secion.

## 4. Peptide strings for protein identification with reduced alphabets

Proteomic analysis is often based on proteolysis to obtain peptides that can be studied more easily than whole proteins. A whole protein is formed by the primary sequence of amino acids folding in various ways to give it secondary structure (helices, sheets, and turns) and tertiary structure from distant residues in secondary structure coming together (such as bonding of two cysteine residues). More often than not this results in many residues in the interior of the structure becoming inaccessible. With the protein broken down into peptides, the structure is reduced to open and more or less linear short chains that are more easily transported in and bound to various media for analysis and whose entire residue sequence is accessible. The proteolytic enzyme used most often for protein digestion is trypsin, a serine endoprotease that cleaves protein sequences on the C-side of Lys (K) and Arg (R) [27]. (In what has been referred to as mirror image behavior the enzyme LysN has a similar effect on the N-side [28].) Much of current protein identification in MS is based on sequencing peptides and locating their parent proteins in a database [29,30]. This approach is used in the Human Proteome Project, the 2023 HUPO Report can be accessed at [31]. As noted earlier it is not necessary to know the full peptide sequence, partial sequences are sufficient [9,26].

Peptides can be viewed as strings on an alphabet. Denote the set of AAs with the alphabet

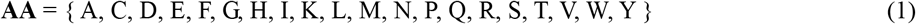

and a character in the alphabet as aa. A peptide can be written as

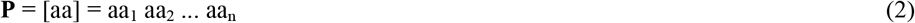

with aa_1_ being the N-terminal residue and aa_n_ the C-terminal residue.

A peptide array is represented by [**P**], where the elements are of variable length | **P** |: Thus

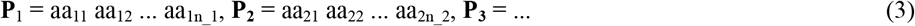

Sequencing a peptide **P**_**i**_ with a carboxypeptidase consists of identifying aa_in_i_ aa_in_i-1_ … aa_i1_ in that order.

Consider peptides obtained by proteolysis of a protein mixture with Lysine at their N-end. There is no need to identify Lysine, as it can only be at the N-end of every peptide. This reduces the alphabet to

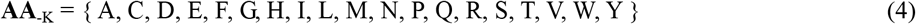

Now consider using carboxypeptidases C, B, and A collectively to identify all 19 AAs. Thus carboxypeptidase C can be used to identify only Proline, carboxypeptidase B to identify only Arginine (although it can identify Arginine or Lysine, the latter is no longer in the alphabet in Equation (4)), and carboxypeptidase A to identify any of the other 17 AAs in

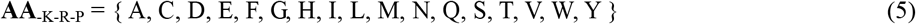

The C-terminal residue of the i-th peptide is identified from the cognate tRNA and is cleaved in one of the following ways:

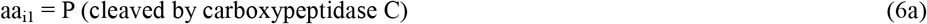

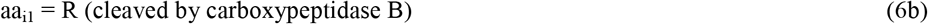

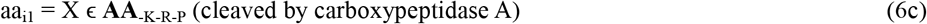

A peptide string obtained by **Procedure** SequencePeptideArrayWithCarboxypeptidaseABC is therefore the reduced string of the form

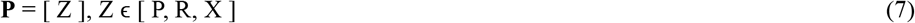

## 5. Protein identification of large peptide arrays cleaved by multiple carboxy exopeptidases: a minimalist approach

C-terminal sequencing has been studied extensively with bulk samples [32-35]. The present study is focused on single molecule sequencing. The following paragraphs describe the use of three carboxypeptidases in single molecule recognition of C-terminal residues of immobilized peptides in succession.

Consider Equation 7. The reduced peptide string for every peptide in the proteome with Lysine at the N-end can be computed and stored along with the identity of its parent protein. Identifying the parent protein of a (reduced) peptide string then consists of comparing it with all the reduced strings for the proteome and outputting the parent (or parents).

The cleavage times for each AA type can be obtained empirically. Let the mean cleavage time for an AA be T_aa_. Alternatively, let the times for the three sets be

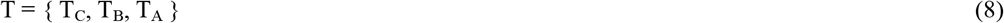

where, for simplicity, it is assumed that cleavage times are the same for all 17 elements in **AA**_-K-R-P_.

If T_C_ = T_B_ = T_A_ = T_Z_ then the cleaving sequence becomes straightforward. Thus after an interval of T_Z_ the C-terminal residues of every peptide in the array will have been cleaved and the next cycle can start. In this case cleaving can be done in parallel by adding all three types to the solution. Thus carboxypeptidase C cleaves all leading Prolines (P), B cleaves all Arginines (R), and A cleaves all the other 17 types of C-terminal residues. This assumes that in every cycle the carboxypeptidase used cleaves the target C-terminal residue without any delayed reaction. That is, the second, third, etc. residues are not cleaved. However if the cleaving times are considerably different there can be skipped residues of any of the three classes, this is discussed below.

A more reliable way of cleaving would be to cleave serially. Thus cleave Prolines first over a time T_C_, Arginines over a time T_B_, then cleave all the others (with carboxypeptidase A). In the last case if some AA_i_ has a cleavage time T_i_ > 2 T_j_ for an AA_j_, j ≠ i, it will result in a skip in the sequence if the next residue is aa_j_. Skip errors are not a major problem as long as | T_i_ - T_j_ | is not too high (otherwise too many residues may be cleaved in delayed reactions over the extended time of T_i_.

Thus the possibility of more than one AA in **AA**_-K-R-P_ being cleaved in one cycle has to be taken into account. The peptides strings that are obtained no longer satisfy Equation 7. Such strings will have the form

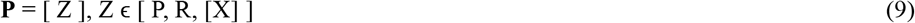

This behavior is taken into consideration in the computational experiments described in the next section, where a random set of peptides obtained from a random selection of proteins from the human proteome is used to obtain protein identification rates under various conditions. The results show that for a large majority of peptides their parent proteins can indeed be identified unambiguously.

As an example, consider protein number 4 with id A6NJR5 and length 290 in the human proteome (Uniprot id UP000005640_9606):

~~~
MQKHYTVAWFLYSAPGVDPSPPCRSLGWKRKKEWSDESEEEPEKELAPEPEETWVVEMLCGLKMKLKQQRVSPILPEHHKDFNSQLAPGVDPSPPHRSFCWKRKREWWDESEESLEEEPRKVLAPEPEEIWVAEMLCGLKMKLKRRRVSLVLPEHHEAFNRLLEDPVIKRFLAWDKDLRVSDKYLLAMVIAYFSRAGLPSWQYQRIHFFLALYLANDMEEDDEDPKQNIFY FLYGKTRSRIPLIALFQKLRFQFFCSMSGRAWVSREELEEIQAYDPEHWVWARDRARLS
~~~

In the reduced alphabet of Equation 5 this becomes

~~~
XXKXXXXXXXXXXXPXXXPXPPXRXXXXKRKKXXXXXXXXXPXKXXXPXPXXXXXXXXXXXXKXKXKXXRXXPXXPX XXKXXXXXXXPXXXPXPPXRXXXXKRKRXXXXXXXXXXXXXPRKXXXPXPXXXXXXXXXXXXKXKXKRRRXXXXXPX XXXXXXRXXXXPXXKRXXXXXKXXRXXXKXXXXXXXXXXXRXXXPXXXXXRXXXXXXXXXXXXXXXXXXXPKXXXXX XXXXKXRXRXPXXXXXXKXRXXXXXXXXXRXXXXRXXXXXXXXXXPXXXXXXRXRXRXX
~~~

Digesting with an endopeptidase that cleaves before Lysine (K) on the N-side results in the following peptides (shown in the reduced alphabet):

~~~
KXXXXXXXXXXXPXXXPXPPXRXXXX KRKKXXXXXXXXXPX KXXXPXPXXXXXXXXXXXX KX KX KXXRXXPXXPXXX KXXXXXXXPXXXPXPPXRXXXX KR KRXXXXXXXXXXXXXPR KXXXPXPXXXXXXXXXXXXKX KX KRRRXXXXXPXXXXXXXRXXXXPXX KRXXXXXKXXRXXX KXXXXXXXXXXXRXXXPXXXXXRXXXXXXXXXXXXXXXXXXXP KXXXXXXXXX KXRXRXPXXXXXX KXRXXXXXXXXXRXXXXRXXXXXXXXXXPXXXXXXRXRXRXX
~~~

or, in abbreviated form, where the exponent is the number of repeats,

~~~
KX^12^PX^3^PXPPXRX^4^ KR K KX^9^PX KX^3^PXPX^12^ KX KX KX^2^R^2^PX^2^PX^3^ KX^7^PX^3^PXPPXRX^4^ KR KRX^13^PR KX3PXPX^13^ KX KX KRRRX^5^PX^6^RX^3^PX^2^ KRX^5^KX^2^RX^3^ KX^11^RX^3^PX^5^RX^19^P KX^9^ KXRXRXPX^6^ KXRX^9^RX^4^RX^10^PX^6^RXRXRX^2^
~~~

They are cleaved from the right to the left by the appropriate carboxypeptidase. For example, the first peptide is cleaved successively by (carboxypeptidase) A, A, A, A, B, A, C, C, A, C, A, A, A, C, A, and A, leaving behind the N-terminal Lysine (K) bound to the slide. Which carboxypeptidase to use is known from the tRNA used in the identification step of a cycle. Thus the sequence of steps is well-defined and unambiguous.

In the next section the protein identified by a peptide is computed for the full human proteome. Following this the identification rate with 1 or 2 residues skipped is computed for a random sample of 1000 proteins. The list of proteins and the set of identifying peptides for every protein in this sample can be found in Supplementary File SI-4.

## 6. Computational results

The proposed model was studied computationally to determine the identification rate for a full proteome and a random sample of 1000 proteins with and without skipping of residues. Sequence data for the human proteome (Uniprot id UP000005640_9606) were downloaded from www.uniprot.org and stored in an abbreviated form containing only the protein id, its length, and the primary sequence. This abbreviated version is given in Supplementary File SI-1. This set was converted to the reduced alphabet {P, R, X} (Equation 7); it can be found in Supplementary File SI-2. All peptides anchored at Lysine (K) at the N-end were extracted; Figure 3 shows their length distribution. (Notice the large number of peptides of length 1 (with the single residue K).) In all there are 653313 peptides in 20598 proteomes, of which 38099 have more than 50 residues.

**Figure 3.**
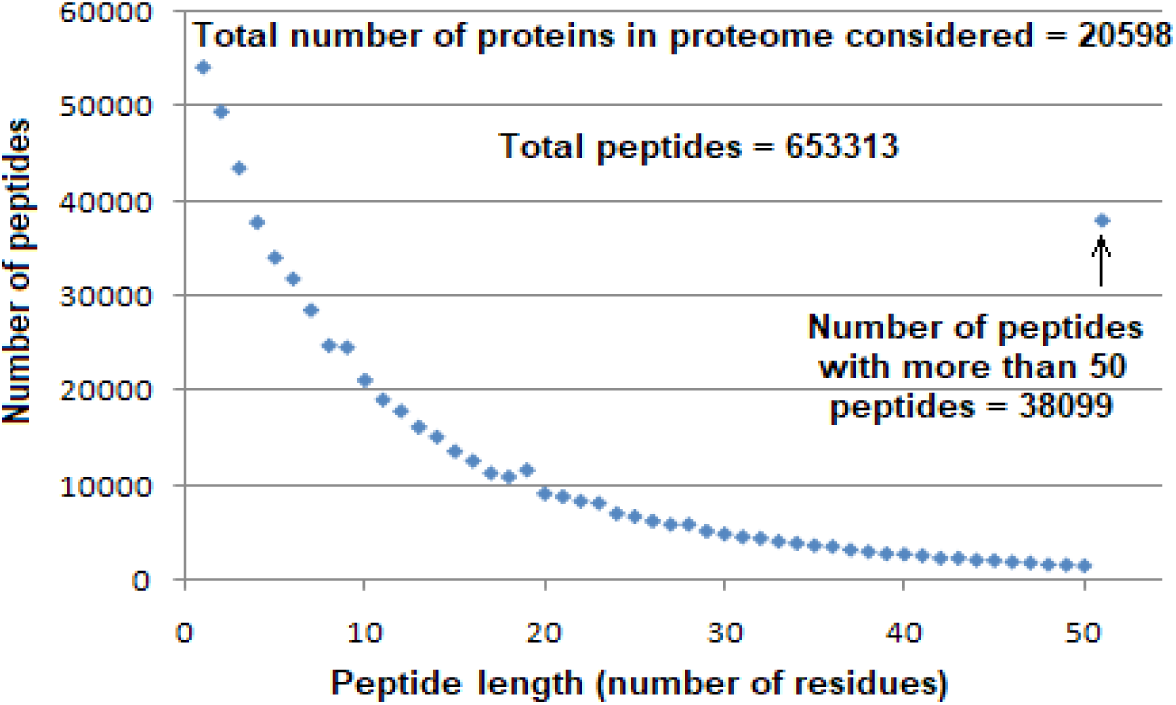
Length distribution of proteolyzed peptides anchored by Lysine (K) at their N-end. There are in all 653313 peptides in 20598 proteomes, of which 38099 have more than 50 residues.

**Figure 4.**
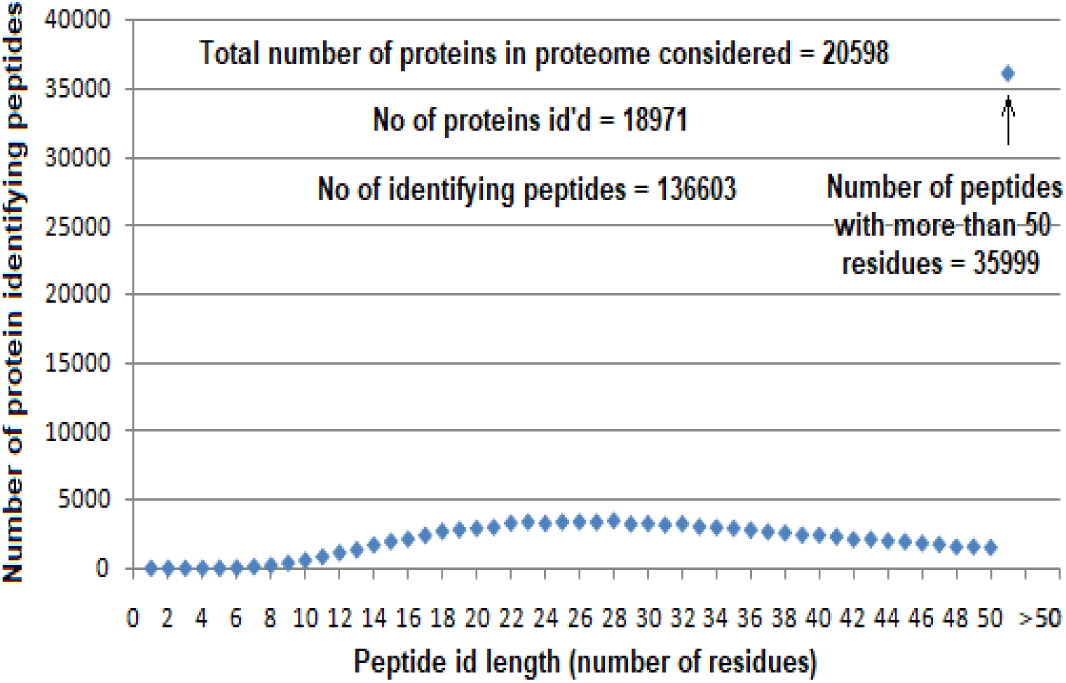
Number of protein-identifying peptides distributed by length. A total of 136603 peptides identify 18971 proteins, of which 35999, or a little over 25%, have more than 50 residues (cf: [26]).

Leaving out peptides with fewer than 10 residues, all other peptides that are ids for their container proteins were computed. Figure 2 shows the length distribution for the number of peptides that are ids. A total of 18971 proteins, or ∼92% of the total number considered have at least one identifying peptide with the reduced alphabet, which shows that partial sequences are sufficient to identify a substantial majority of the proteins in the proteom.

To determine the effect of skipped residues due to delayed cleaving actions of carboxypeptidases (A, B, and C), a random sample of proteins from UP000005640_9606 was generated; a list containing their numbers in serial order from the proteome can be found in Supplementary File SI-4. First the identifying peptides without any skipping of residues was computed; following this the first and second X residue in every peptide were skipped one after the other and the identification rates were computed in each case. Figure 5 shows the results in the three cases. Skipping of residues does not seem to have a significant adverse effect on the identification rate. However further computations may be required before a definitive conclusion can be drawn on this.

**Figure 5.**
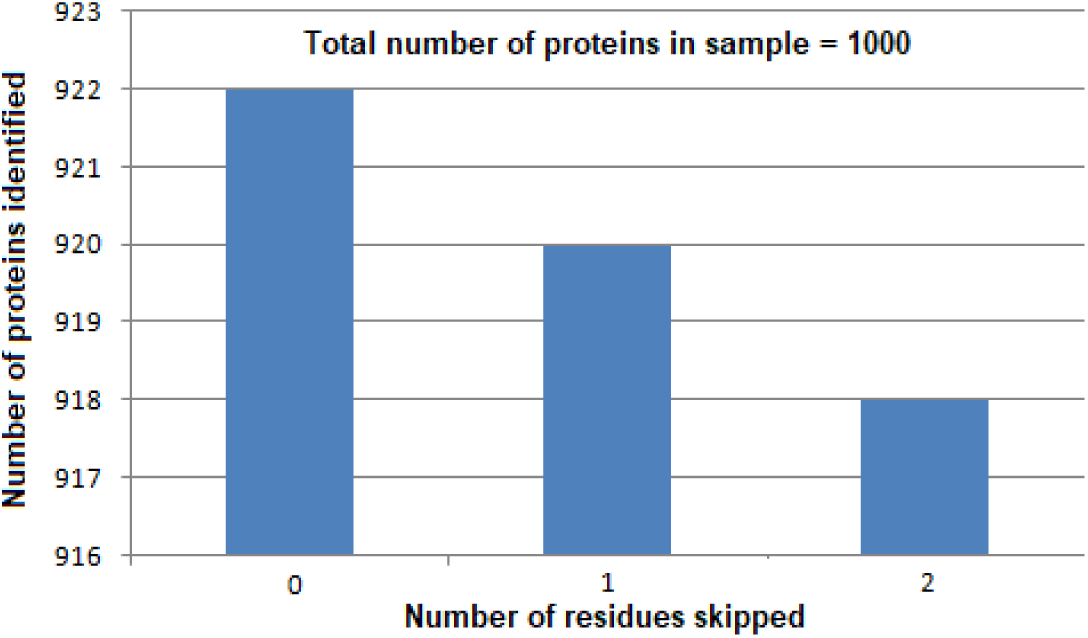
Protein identification rates with and without skipping of residues. With 1000 sample proteins, more than 92% are identified without residues being skipped. With one or two residues in every peptide in every sample protein reduction in identification rate is negligible.

## 7. Extension to full peptide sequencing and whole protein sequencing

Leaving out Lysine (K), as it is always the N-end residue of a peptide, let the empirically obtained mean cleavage times for the other 19 AA types be given by the set

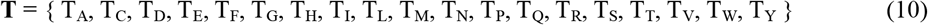

Since the three carboxypeptidases are mutually exclusive in their cleaving actions, T can be partitioned into three classes:

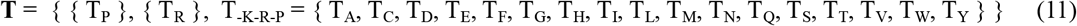

Because of exclusivity the cleaving times of P and R do not affect each other or those of the other 17 in T_-K-R-P_, and the identification and cleaving cycles of P and R are independent of each other and of the AAs in T_-K-R-P_. If the cleaving times in T_-K-R-P_ are different, residues may be skipped, as noted earlier, resulting in partial sequences obtained. The cleaving times of P and R are independent of the AAs in A_-K-R-P_.

Now consider ordering the cleaving times in T_-K-R-P_ in increasing order:

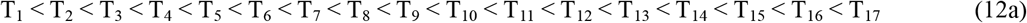

where the subscripts in the ordering are mapped 1-1 to the elements in T_-K-R-P_; and P and R similarly as

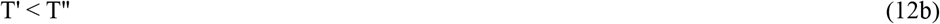

If recognition of terminal residues and subsequent cleavage are done such that this order is preserved across the breadth and depth of the peptide array then there will be no skipping of residues and full sequences can be obtained for all the peptides in the array. Recognition-cleavage has a recursive structure because the next terminal residues exposed after the currently cleaved terminal residues peptides may be any one of the 19 AA types. The formal procedure to do this is given next (T_P_ < T_R_ is assumed).

**Procedure** SequencePeptideArrayWithoutSkippingResidues

**while** there are no more peptides to sequence **do**

**do**

Identify Proline terminals, cleave all Prolines with carbopeptidase C

**until** there are no more Proline terminals

**do**

Identify Arginine terminals, cleave all Arginines with carbopeptidase B

**until** there are no more Arginine terminals

**for** i=1 **to** 17 **do**

**do**

Identify AA_i_ terminals, cleave all AA_i_s with carbopeptidase A

**until** there are no more AA_i_ terminals

**call** SequencePeptideArrayWithoutSkippingResidues

**end for**

**end while**

On exit from the above procedure all the peptides will have been fully sequenced.

Although this may appear to be a tedious and time-consuming process, most of the steps are part of standard practice so it should be possible to automate the protocols.

### Complexity of skip-less sequencing

The worst-case complexity occurs when exactly one terminal residue is cleaved in each call. But this is not of concern because real protein sequences do not have this extreme structure. On the average about half the terminal residues will be cleaved in a single cycle, so the complexity of the recursive procedure will be a low degree polynomial of M.

In the above development implicitly it has been assumed that T_i+1_ - T_i_ in Equation 12 is ∼T_i_. If T_i+1_ - T_i_ is small then the two cleavages can occur closely in sequence and may have to be done in the same cycle. This can be done by using two different colors for the fluorescent dyes. Imaging systems usually have a level of resolution that can accommodate recognition of up to 4 colors (as illustrated in next generation DNA sequencing), so there is some leeway available here.

## 8. A minimalist procedure for amino acid identification in the bulk

The AA identification method given above can be adapted for implementation in the bulk with a minimal number of steps using only equipment available in an undergraduate chemistry lab. Most of the steps involved are much simpler than the ones in Section 2 and can be carried out with a centrifuge and pipettes for separation of reactants and by detecting the presence of AA in the supernatant with the ninhydrin test. Separation steps transfer fractional materials to designated receptacles for collection and reuse; thus expensive reagents like AARSs and tRNAs can be reused either with repeat experiments or with successive AAs to fully sequence a peptide, or partially using only three carboxypeptidases to obtain a string such as in Equation 9 for identification of a protein in a database.

The following is an outline of the procedure, with buffers and other additives added as required. It assumes use of a centrifuge that can hold N ≥ 20 test tubes; if N < 20 the procedure may be run multiple times depending on N. Following usual laboratory practice, the active test tubes are to be distributed for rotor balancing with dummy test tubes containing appropriate levels of inactive solution with the required volume or weight.

**Procedure** AminoAcidIdentificationInTheBulk

Input: Amino acid aa_unknown_

1. **For** i = 1 to 20 **do** the following:
  1a. Add aa_unknown_ with AARS_i_, tRNA_i_, and ATP in appropriate amounts to test-tube_i_ along with suitable buffer
  1b. Shake, stir, or otherwise aid reaction among reactants in test-tube over desired time period
  1c. Load test-tube_i_ onto centrifuge
2. Load-balance centrifuge as required
3. Perform centrifugation with appropriate speed for suitable length of time
4. **For** i = 1 to 20 **do** the following:
  4a. Pipette out supernatant_i_ from test-tube_i_ into labeled-container_i_ while leaving sediment at bottom of test-tube_i_
  4b. Add NaOH to sediment in test-tube_i_
  4c. Shake, stir, or otherwise aid reaction among reactants in sediment over desired time period
5. Load-balance centrifuge as required
6. Perform centrifugation with appropriate speed for suitable length of time
7. **For** i = 1 to 20 **do** the following:
  7a. Add suitable amount of ninhydrin to supernatant_i_ in test-tube_i_
  7b. If supernatant turns blue/violet, output aa_unknown_ = aa_i_
  7c. Pipette out supernatant

Notes A:

1a. Amounts to be determined empirically.

1b. Time period to be determined empirically.

3. Time period to be determined empirically.

4a. Contents of container i ≠ j should include uncharged aa_i_ and unreacted ATP; contents of container i = j should include excess (uncharged) aa_j_, excess unreacted ATP, AMP from charging of tRNA_j_ with aa_j_; sediment in test-tube_i≠j_ should contain AARS_i≠j_, (uncharged) tRNA_i≠j_; sediment in test-tube_i=j_ should contain AARS_i=j_, amino-acyl_i=j_-tRNA_i=j_, excess (uncharged) tRNA_i=j_.

4b. Amount to be determined empirically.

Notes B:

1. Alternatively, supernatant_i_ in Step 4a can be examined for the presence of AMP; if it is present aa_unknown_ = aa_i_ can be output and the loop can be exited.
2. Similarly the loop in Step 7 can be prematurely exited as soon as the ninhydrin test shows positive, provided there are no errors or any anomalous behavior displayed by the reactants.
3. Following Equation 6, the above procedure can be used in truncated form (with N = 3) to identify the input aa_unknown_ as P (Proline), R (Arginine), or X (other).

### Efficacy analysis

1. Cost: A downside to the proposed method, whether at the SM level or in the bulk, is the high cost of reagents and the general difficulty in their procurement. Additionally, availability is usually not immediate and involves considerably delays in delivery as noted in publicly posted information from providers on the web. Prices posted are in the neighborhood of $400-450 for 1 mg of an AARS. The price of a specific tRNA is almost double at $850/mg, while tRNA mixtures extracted from *E. coli* cost considerably less; this large difference is primarily because tRNA purification is a complex and time-intensive process. However this situation can be alleviated by noting that tRNA charging requires the cognate AARS to be present. In the setups described above (both SM and bulk), since each AARS is in a different cavity or test-tube, the tRNA used does not have to be pure-even with a tRNA mixture the correct result can be obtained if it is assumed that mischarging does not occur. The cost issue in the case of AARS can be mitigated by reuse, as noted in Item 2 next; the time factor is addressed in Item 3 below.
2. Recovery and reuse: Note that following AA detection AARS_i_ and tRNA_i_ remain in the sediment in test-tube_i_ and can be reused as is. This is because AARS_i_ and AARS_k_ i ≠ k, are never together. Similarly, if pure tRNA is used, tRNA_i_ and tRNA_k_ i ≠ k, are never together; however as noted in Item 1 above, a tRNA mixture can be used without adverse effects.
3. Synthesizing reagents for in-house use: One way to minimize explicit costs is to make the required reagents in-house. Pure tRNAs can be synthesized chemically (using the phosphoramidite method) or enzymatically (using the TdT enzyme) in the lab. Chemical oligo synthesis, which is typically limited to about 100 nucleotides before secondary structure issues get in the way, are not an issue as all the known tRNAs have around 75-90 nucleotides. The downside is the toxic waste generated. While enzymatic synthesis is generally water-based the issue of enzyme cost becomes an inhibiting factor. The real limiting factor then is the availability of AARSs. It is not easy to synthesize them, as they are about 300-500 residues long, and peptide synthesis is a complex process that generates unacceptably large amounts of toxic waste. While emerging peptide synthesis methods that are more environmentally acceptable continue to be the focus of ongoing studies, developing AARSs in-house is not a viable option.

## 9. Discussion

1. The minimalism in the proposed approach is threefold: the number of tRNAs, the number of carboxypeptdidases, and the number of steps used in a cycle. Thus at most one or two tRNA types (tRNA_P_ and tRNA_R_) are required (and therefore only one or two cognate AARS types). Notice that none of the other 18 are required. Only 3 carboxypeptidases are required; and the number of steps in a cycle is limited to 4: tRNA charging, imaging, tRNA dissociation, and terminal residue cleaving. The last two steps can be combined into a single step by cleaving the reside with the tRNA still charged, if it is assumed that the presence of the tRNA does not adversely affect the cleavage reaction. Notice that this also allows determining the end of the cleavage step because the latter will be indicated by the absence of fluorescence at the terminals of the affected peptides. (The method also suggests a straightforward way to determine cleavage times for the 20 AAs.)
2. The method is not biased toward any protein, so the extent to which the dynamic range problem is addressed and resolved is determined entirely by the distribution of component proteins in the sample.
3. The method may be considered to have a high green coefficient as almost all reagents used can be recovered from the wash and reused. It may be noted in this context that the identity of the reagent is known from the step in the cycle after which washing is done; this can be used for routing a recovered reagent to its designated receptacle.
4. With the use of a single type of protease the resulting peptides often tend to be long and complex. This has led to the use of multiple proteases, sometimes referred to as multidimensional proteolysis [36-38]. In particular, there has been a move toward alternatives or complements to trypsin, which has dominated protein sequencing, especially with MS [39-40]. With the use of multiple proteases, the number of residues to consider is reduced. This can be used to advantage in the method proposed here as it alleviates the delayed cleaving problem. See [41] for a summary of different types of proteases, and [42] for the statistical properties of the peptides that result from cleavage of multiple AA types.
5. One of the hallmarks of Next Generation Sequencing of DNA is that the sequences of whole chromosomes can be obtained by shotgun sequencing of overlapping subsequences and associated algorithms to construct the full sequence. Similar methods can be used to obtain the primary sequences of whole proteins from the sequences of overlapping peptides obtained through the use of multiple proteases, for example peptides with N-terminal Arginine (R), Aspartic Acid (E), Glutamic Acid (D), etc. In particular a protein can be broken into peptides with desired start and end residues. For example LysN creates peptides starting with K, while formic acid creates peptides ending in D. When these two are applied in sequence (in any order), the result is peptides that start with K and end with D. With four peptidases/chemical agents, there are 24 different orders of application; see [41,42]. The optimum order may be determined experimentally.
6. The bulk procedure described above has a minimal number of steps and uses equipment found in a college chemistry lab. As such it is well-suited to student experiments that can be conducted using skills acquired early in an undergraduate program. The notes in Section 8 suggest that there is considerable scope for a variety of empirical studies based on the procedure given here, which can serve as a platform for a multitude of junior/senior-level projects.

## Supporting information

human proteome sequence

SI-1 with reduced alphabet

peptides from SI-2 anchored at Lysine

random proteins and their identifiers

## Supplementary Information Files

SI-1: human proteome sequence information (Uniprot id UP000005640_9606) in abbreviated form; SI-2: SI-1 with a reduced alphabet; SI-3: all peptides from SI-2 anchored at Lysine (K) at the N-end; SI-4: 1000 randomly selected proteins from human proteome by Uniprot id and their identifying peptides without any skipped residues.

